# 3D-printed drug testing platform based on a 3D model of aged human skeletal muscle

**DOI:** 10.1101/2020.06.18.158659

**Authors:** Rafael Mestre, Nerea García, Tania Patiño, Maria Guix, Judith Fuentes, Mauricio Valerio-Santiago, Núria Almiñana, Samuel Sánchez

## Abstract

Three-dimensional engineering of skeletal muscle is becoming increasingly relevant for tissue engineering, disease modeling and bio-hybrid robotics, where flexible, versatile and multidisciplinary approaches for the evaluation of tissue differentiation, functionality and force measurement are required. This works presents a 3D-printed platform of bioengineered human skeletal muscle which can efficiently model the three-dimensional structure of native tissue, while providing information about force generation and contraction profiles. Proper differentiation and maturation of myocytes is demonstrated by the expression of key myo-proteins using immunocytochemistry and analyzed by confocal microscopy, and the functionality assessed *via* electrical stimulation and analysis of contraction kinetics. To validate the flexibility of this platform for complex tissue modelling, the bioengineered muscle is treated with tumor necrosis factor α to mimic the conditions of aging, which is supported by morphological and functional changes. Moreover, as a proof of concept, the effects of Argireline^®^ Amplified peptide, a cosmetic ingredient that causes muscle relaxation, are evaluated in both healthy and aged tissue models. Therefore, the results demonstrate that this 3D-bioengineered human muscle platform could be used to assess morphological and functional changes in the aging process of muscular tissue with potential applications in biomedicine, cosmetics and bio-hybrid robotics.

## 1. Introduction

Recent developments in 3D bioprinting and tissue engineering have opened the possibility of mimicking native tissue by the biofabrication of engineered three-dimensional muscle models that can mimic the extracellular environment of native tissue^1–4^, offering opportunities for studies in muscle development or muscular diseases, thus avoiding the need for animal models that can carry ethical issues. Moreover, the integration of skeletal muscle tissue with artificial materials has led to a wide variety of innovative applications in the field of bio-hybrid robotics and bio-actuators^5^, with examples of miniaturized bio-robots that can crawl^6,7^ or perform simple forms of actuation^8^. In particular, bio-actuators, born as a starting point towards complex bio-robots, have become excellent candidates for biosensing and drug screening platforms, providing information about contraction kinetics of three-dimensional muscle tissue, as well as their maturation and their adaptability^2,9^.

Muscle-based bio-actuators usually consist of a 3D bio-engineered skeletal muscle strip or ring surrounded by two or more cantilevers that provide mechanical support to assist in the differentiation and maturation of these cells^10^. Different models employ cantilevers that mimic muscular structures, using their deflection to give estimates of the forces generated during tissue formation and compaction, as well by active contractions^2^. Electrical or optical stimulation of 3D-bioengineered muscle tissues can give insightful information about the contraction dynamics and the generation of force^11^, which can then be used to analyze how specific drugs affect these parameters^12,13^, and provide useful data for tissue engineering and, in particular, for the fabrication of relevant skeletal tissue models.

Both forces and contraction kinetics can provide valuable information in the study of muscle functionality. For instance, the range of myopathies that can affect the function of skeletal muscle in the form of muscle weakening can be very heterogeneous and with a variety of aetiologies. Consequently, there is a need for human tissue models to perform efficient screenings of a wide variety of treatments by measuring physiological parameters, such as force and contraction kinetics of diseased tissue. Although two-dimensional skeletal muscle cultures are widely known and have been extensively studied, they lack 3D architecture, are difficult to maintain in the long-term and measuring their forces is challenging. Moreover, more complex three-dimensional models of skeletal muscle tissue have been mainly focused on murine primary cells or cell lines, with increasing applications in the hybrid bio-robotics field^2,6,7,14^, although some cases have been dedicated to pharmacological testing or biomolecule monitoring^4,13^. Nonetheless, bio-engineering of human muscle cells has received some attention, yet most of the studies have primarily aimed at studying the effects of mechanical stretching^15^, cycling^16^ or cell density^15^. In this regard, related to the study of forces in human skeletal muscle tissue, Madden et al. successfully developed a human myobundle platform from myoblasts, focusing on obtaining clinically relevant responses and analyzing the tetanic and twitch behaviors of their models^17^.

Studies on cell aging have found an interest in skeletal muscle research, since a deep understanding of the aging process of skeletal muscle can provide valuable information about tissue functionality and its relationship with the increasing number of myopathies that the elderly can suffer. Tumor necrosis factor α (TNF-α) is a cytokine that has been described as one of the main drivers for the known morphological changes that take place in aging muscles^18–20^, and has been linked to the development of sarcopenia affecting satellite cells^21^. In mouse cell lines, differences in TNF-α have been related to changes in calcium transients^22,23^ and reduction of contractile force^20^. Furthermore, *in vitro*, this cytokine is known to have an effect on myogenesis inhibition^24^ and cause atrophy^25^, as demonstrated in C2C12 cell lines, as well as having an important effect in muscle regeneration after injury *in vivo*^26^. Sarcopenia has also been linked to the presence of macrophage-secreted TNF-α in men during aging^32^. Several cellular pathways have been associated with elevated TNF-α activity in aged muscle, such as reactive oxygen species^18,24^, the sphingolipid metabolism^19^ or autophagic degradation^27^, with the up- or downregulation of several genes as drivers.

The study of aging in skeletal muscle tissue has also relevant applications in the cosmetics field, since skin manifestations of aging have been linked to physiological and functional changes of subcutaneous tissues, such as adipose tissue and skeletal facial muscle^28,29^. In particular, expression wrinkles, caused by repetitive muscle contractions, have been traditionally targeted by muscle relaxants, like Botox injections^30,31^. This mechanism of action relies on the inhibition of neurotransmitter release by motor neurons, which reduce the frequency of facial muscle contractions^31,32^. Muscle aging is associated with an increase of muscle relaxation time^33–35^. In fact, it was described that facial muscles increase their resting tone in old subjects contributing to the appearance of structural aging features in the elderly.^36^ Focusing on this concept, the enhancement of a facial muscle relaxed state, working on a post-synaptic approach, may contribute to minimize expression wrinkles in old people. In addition, a partial reduction of maximal force exerted by facial muscles could reduce the deformation of the skin layers located above of these muscles, leading to a visual reduction of the expression wrinkles. Hence, by treating healthy human myocytes with TNF-α, it might be possible to mimic the morphological and functional characteristics of aged muscle, like decreased force and increase in muscle relaxation time. Moreover, applying high frequency stimulations could simulate the sustained contractile state present in expression wrinkles and the effects of cosmetic ingredients acting to reduce them on a post-synaptic level assessed.

Herein, we present a 3D-printing-based platform for human muscle tissue engineering and drug testing in aged muscle models, where we analyze the contraction kinetics upon low and high frequency stimulations. We demonstrate that 3D-differentiated myocytes show contractile response upon electrical pulse stimulation and contraction patterns similar to those of native tissue. We validate the capabilities of this drug testing platform by the treatment of human tissue constructs with TNF-α, thus obtaining a model of skeletal muscle tissue affected by aging, characterized by atrophy, a decrease in force generation, fiber diameter and disruption of cell sarcolemma. Moreover, as a proof-of-concept, we verify the 3D-printed testing platform by evaluating the efficacy of a green-chemistry-based cosmetic ingredient in development, Argireline^®^ Amplified peptide, an evolution of the commercially available Argireline^®^ peptide with relaxing effects with a pre-synaptic mechanism of action^37,38^. In this study, we evaluate its impact in both aged and non-aged 3D muscle models, focusing only on the post-synaptic component of the neuron-muscular junction. This novel approach allows us to test its relaxing effects after electrical-stimulation-induced muscle contractions. Overall, these results demonstrate the versatility of this human 3D muscle model platform for the study of muscle functionality of aging-related myopathies induced by TNF-α, as well as the opportunities for drug testing in the biomedical or cosmetic fields.

## 2. Results and discussion

Human skeletal muscle tissues (hSMTs) were bioengineered by embedding human myoblasts in a mixture of biopolymers mimicking the ECM found in native tissue. The mixture consisted of Matrigel^®^ and fibrinogen as the main components, as already reported elsewhere for murine skeletal muscle cells^14^. The collagen contained within Matrigel^®^, as well as its other components, helped recreate the collagen- and laminin-rich environment of the basement membrane of the sarcolemma of skeletal muscle tissue^39^. Fibrinogen, also rich in cell-attachment motifs, helped modulating the final stiffness of the matrix (after crosslinking with thrombin) to achieve the right mechanical properties for cell survival and tissue manipulation (**Figure 1**)^2^.

**Figure 1.**
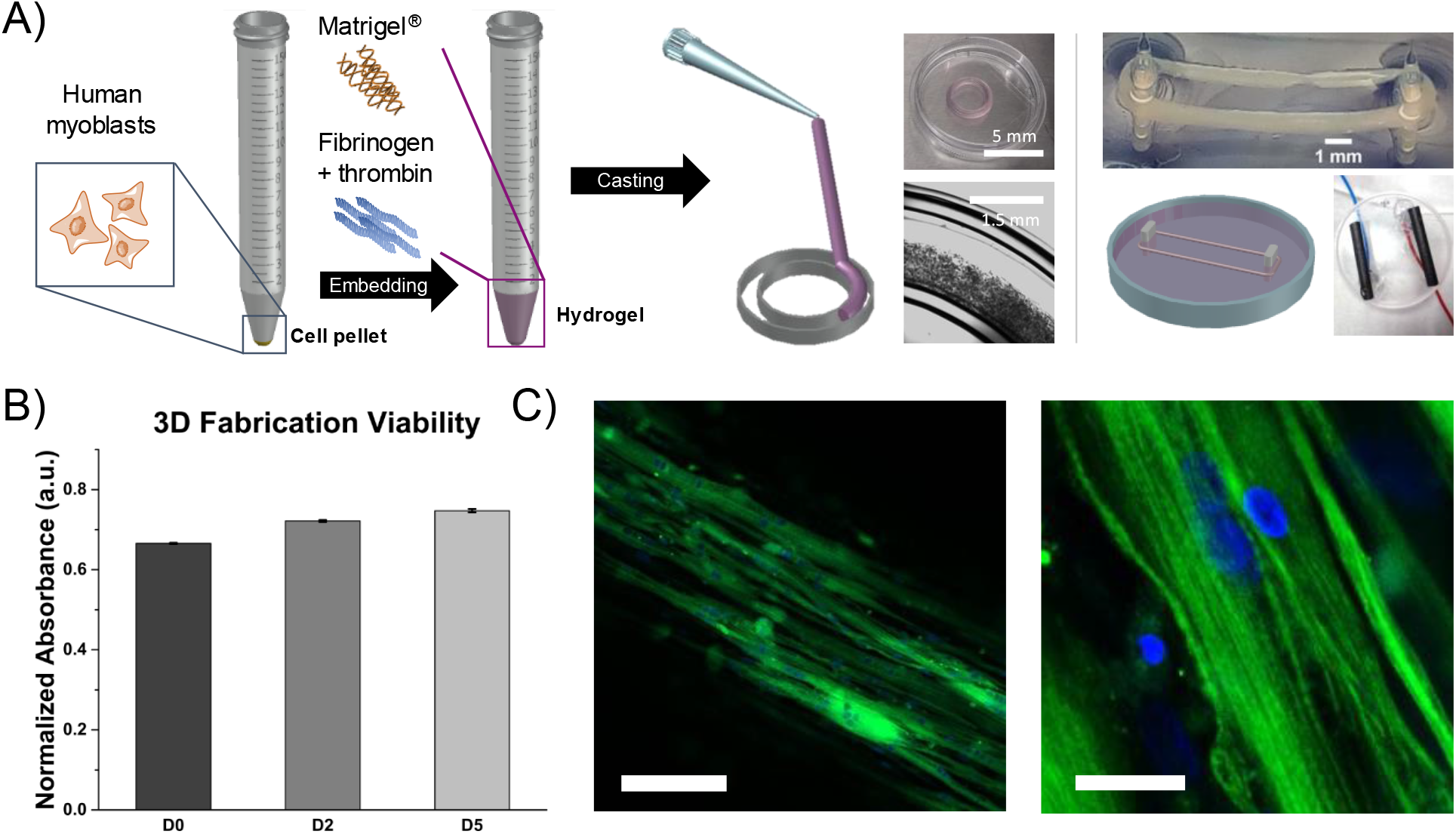
Fabrication of three-dimensional human skeletal muscle constructs. A) Schematic of the fabrication method, based on casting on 3D-printed molds and subsequent transfer into 3D-printed PDMS posts, which is then stimulated with a home-made setup of carbon-based electrodes. B) Metabolic activity PrestoBlue™ assay of the hSMCs several days after differentiation (*N* = 4; mean ± standard error of the mean). C) Immunostaining of MyHC (green) and cell nuclei (blue) after 10 days of differentiation showing well aligned myotubes and (zoom) sarcomeric structure. Scale bars: 125 μm (left) and 20 μm (right).

Human myoblasts were embedded with the hydrogel mixture (Matrigel^®^, growth medium and thrombin) at a final concentration of 5 million cells/mL. Thereafter, fibrinogen was added, mixed thoroughly, and 80 μL of the solution was casted in a 3D-printed mold made of polydimethylsiloxane (PDMS), rapidly in order to avoid the crosslinking of fibrinogen into fibrin induced by thrombin. Afterwards, the cell-laden hydrogels were left in an incubator for Matrigel^®^ to fully crosslink. After 30 min, growth medium (GM) supplemented with 6-aminocaproic acid (ACA) was added and maintained for 2 days. The addition of the anti-fibrinolytic agent ACA was a crucial step in order to obtain hSMTs that were not affected by the strong degradation of collagen and fibrin by plasmin, cysteine cathepsins and other proteases^40^. We found that, after 5 days of differentiation, the tissue would completely degrade due to the action of these proteases. Inhibition of serine protease by the addition of ACA ensured structural integrity for the duration of all our studies. Beforehand, we performed studies of cell growth, proliferation and differentiation of human myoblasts in 2D, which showed no differences when the media were supplemented with ACA. Moreover, previous reports in the literature demonstrate beneficial aspects of using ACA to increase the lifetime of bio-actuators without loss of function^40^, showing that addition of 1 mg/mL of ACA can increase the life expectancy of murine muscle tissue constructs from 8.2 to 17 days.

After 2 days in GM, the hSMTs were carefully transferred to a 3D-printed two-post system and the media changed to differentiation media (DM; both media were always supplemented with ACA, even if not explicitly stated). Tissue compaction after transfer into the two-post system was minimal. This differs particularly from previous reports using murine skeletal muscle cells (C2C12 cell line), in which compaction of the cell-laden hydrogel was substantial^2^. Since static mechanical stimulation of myocytes is crucial for proper differentiation and necessary for post deflection and force measurement, the dimensions of the molds had to be carefully designed for the tissue to perfectly adapt to the two-post system. Moreover, on the first day after casting, the tissues were gently separated from the outer wall of the molds, due to the high adhesion of the hydrogel to PDMS. In this way, on the second day, before transfer, some compaction of the tissue was already observed (Figure 1A, center). We hypothesize that these human myoblasts seem to be less active than their murine counterparts, perhaps due to their larger size, something also supported by the low release of cells from the scaffold into the bottom of the dish, a very common event with C2C12 cells.

Biocompatibility and cell survival was checked with a metabolic activity PrestoBlue*™* assay (Figure 1B). Several days after differentiation, the normalized absorbance of the samples increased slightly, indicating an increased metabolic activity, and therefore demonstrating the viability of the cell-laden hydrogel during the differentiation process. The optimal differentiation and maturation of the myocytes was further checked by immunostaining of Myosin Heavy Chain (MyHC) and nuclei, which showed elongated and aligned myocytes (Figure 1C). Sarcomeric structures could also be observed under high magnification, indicating maturation of the tissue. Spontaneous contractions were not observed during differentiation.

Maturation of the hSMT was further confirmed *via* electrical stimulation (**Figure 2**). By using a home-made setup of carbon-rod electrodes, we performed electrical stimulation of the hSMTs in the two-post platform after 6 days of differentiation. As expected, the tissues could contract following the applied frequency, as can be observed in Figure 2A and Supplementary Video S1, after a 1 Hz stimulation that produced twitch contractions. Moreover, proper biochemical response was also verified by calcium imaging under fluorescence microscopy, as Supplementary Video S3 and Supplementary Figure S1 show. Other behaviors related to different frequency profiles were also investigated. Figure 2B shows the contractile response after a frequency sweep from 5 Hz to 150 Hz in 5 s intervals. A behavior similar to that of wave summation in native tissue can be observed^33^. However, for higher frequencies (50 Hz and beyond), this response disappears, and a steady tetanic state is maintained until stimulation is stopped. In order to get a grasp on this high frequency response, we examined the contraction kinetics under a high frequency stimulation of 75 Hz for 55 s (Figure 2C). When stimulation starts, there is an initial contraction of high force, which gradually decreases and remains in a tetanic state until stimulation stops. In general, we observed this kind of behavior for all frequencies in the testing range of 10-75 Hz, as well as from day 5 to 12 of differentiation, when our experiments took place. Moreover, this behavior differs from that observed in 3D-bioengineered C2C12 cells, which only show a sustained tetanic contraction for frequencies higher than 10 Hz without an initial stroke^2^.

**Figure 2.**
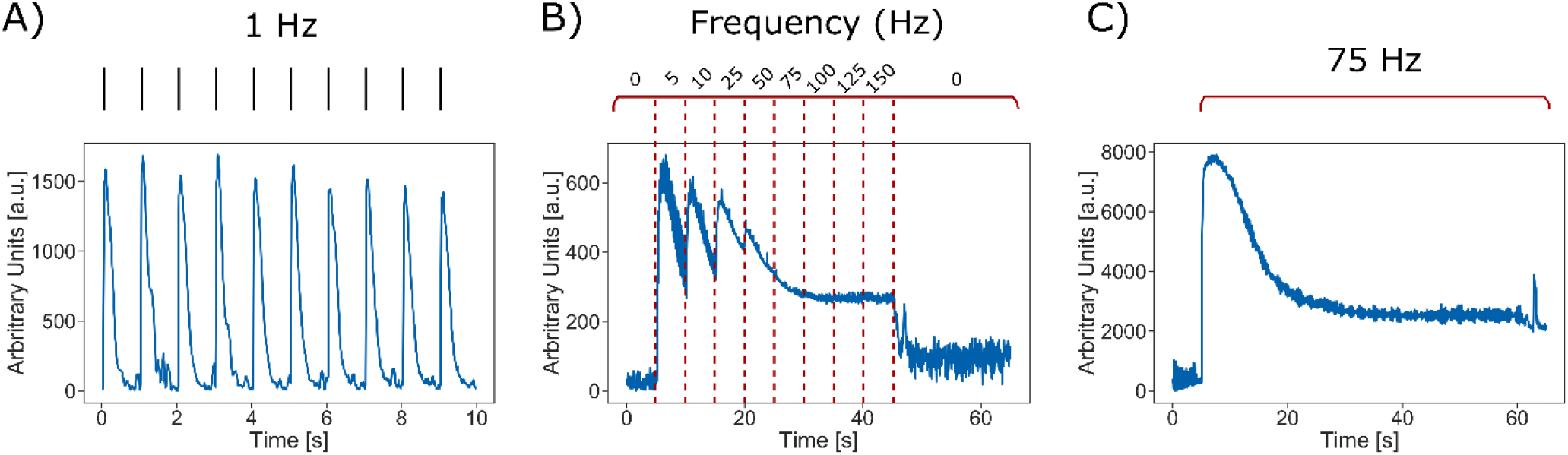
Response of hSMT constructs upon different types of electrical stimulation, mimicking the behavior of native skeletal muscle tissue, at day 6 of differentiation. A) A constant stimulation at equally spaced intervals (1 Hz) produces single-twitch contractions, where the muscle has enough time to fully relax between contractions. B) An increasing frequency sweep in intervals of 5 s exhibits the effect of wave summation, where one stimulation wave builds up over the next one, not allowing the muscle to relax. It is interesting to notice how, for a single frequency, there is a fast decrease in the contraction. C) This effect can be clearly observed in a sustained high-frequency contraction of 75 Hz for 55 s, where the muscle achieves a peak of maximum contraction, followed by a slow decrease, remaining at a baseline level.

TNF-α is a proinflammatory cytokine known to be related to sarcopenia, or loss of muscle mass and function in ageing, as well as reduction of cell fusion^20–26^. Hence, we examined whether addition of this cytokine could be used to model aged skeletal muscle tissue in our muscle platform. We evaluated three different concentrations of TNF-α in the culture media: 20 ng/mL, 40 ng/mL and 80 ng/mL. Morphological and physiological changes were assessed by immunostaining of MHC and cell nuclei 24 h after the addition of TNF-α (**Figure 3**). All concentrations showed multinucleated and aligned myocytes, although morphological differences could be observed with higher concentrations (Figure 3A). White arrows indicate areas where the myocytes show thinning of the fibers with uncharacteristic distortions. Besides, there was a decrease in myocyte diameter, going from more than 30 μm for control (untreated) samples to less than 20 μm for the highest concentration (Figure 3B). Nucleation was also affected by the cytokine, since the number of nuclei per visible fiber was reduced from 9 to 5 for the concentration of 80 ng/mL (Figure 3C). All these morphological changes were consistent with those related to sarcopenia induced by TNF-α^20,21,25,41,42^. In order to test whether functional changes were also compatible, we set the concentration of TNF-α to 40 ng/mL for the remaining experiments, since it was the concentration for which some effects could be observed without becoming too adverse.

**Figure 3.**
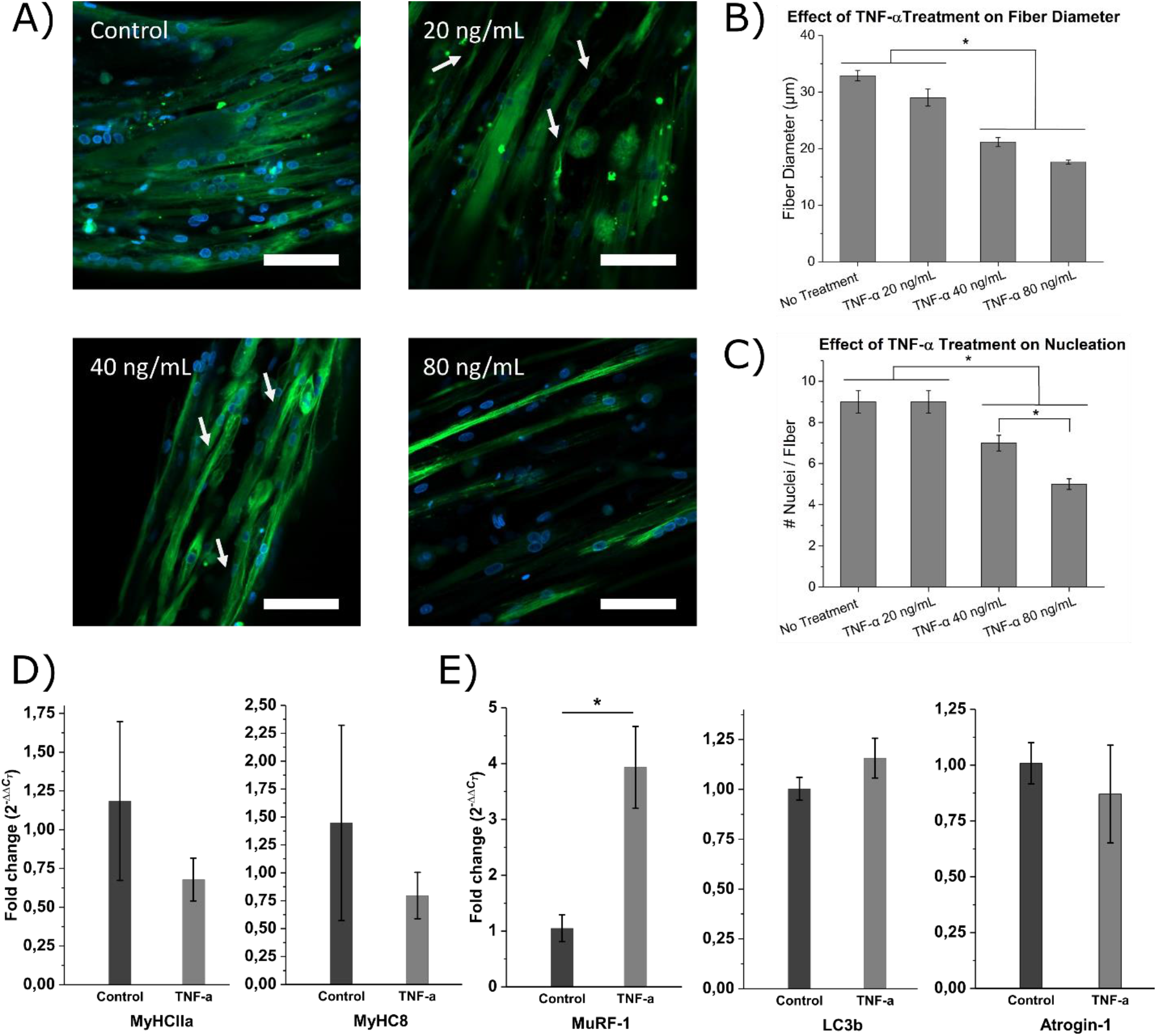
Effect of the addition of Tumor Necrosis Factor α (TNF-α) to three-dimensional hSMC constructs. A) Upon the addition of higher concentrations of TNF-α, the myotubes seem to show a significant decrease in their width and shape distortions (white arrows). MyHC (green) and cell nuclei (blue). Scale bars: 80 μm. B) The diameter of the myotubes decreases with increasing concentrations of TNF-α (N=12; mean ± standard error of the mean). * *p* < 0.05 (Tukey’s HSD test). C) The number of nuclei per fiber is also affected by the concentration of TNF-α (N=12; mean ± standard error of the mean). * *p* < 0.05 (Tukey’s HSD test). D) E) RT-qPCR of the expression of several genes of control samples (untreated) and samples treated with TNF-α at 40 ng/mL for 24 h (N = 3-4; mean ± standard error of the mean). * *p* < 0.05 (Student’s t-test).

The expression of several myosin heavy chain isoforms (namely MyHCI type 1X, MyHCII type 2A and MyHC8 neonatal type) was evaluated by real time quantitative polymerase chain reaction (RT-qPCR) to elucidate whether treatment with 40 ng/mL of TNF-α for 24 h was affecting their expression (Figure 3D). MyHCII and MyHC8 were slightly downregulated for TNF-α-treated samples with respect to control samples, although with high variability, which could be related to the atrophy previously observed by immunofluorescence. Likewise, we studied the expression of several genes that have been linked to atrophy and sarcopenia caused by TNF-α (Figure 3E). MuRF-1, which has been correlated to loss of muscle function in mice^20^ and atrophy in C2C12 cells^25^, was robustly upregulated with respect to untreated tissues. Likewise, the expression of LC3b, a marker of autophagy, has also been shown to increase after addition of TNF-α,^27^ and was slightly upregulated in our experiments, although without statistically significant differences. Atrogin-1, a gene linked to the process of atrophy and, in particular, to ROS-mediated signaling induced by TNF-α^18^, did not show any changes in expression. These results recapitulate the complexity of aging and atrophy in muscle tissue, in which different signaling pathways can be linked to the process in different manners.

Having obtained a morphologically relevant model of native and aged hSMT, we proceeded to use this system as a force measurement platform for drug testing (**Figure 4**). The contractions of the skeletal muscle induced by electrical stimulation could deflect the two PDMS posts, as depicted in Figure 4A. This displacement could then be translated into force generation^2^, as described in the methods. In this way, we could not only assess the kinetics of the contraction, but also the total force generated by the muscle in a quantitative manner.

**Figure 4.**
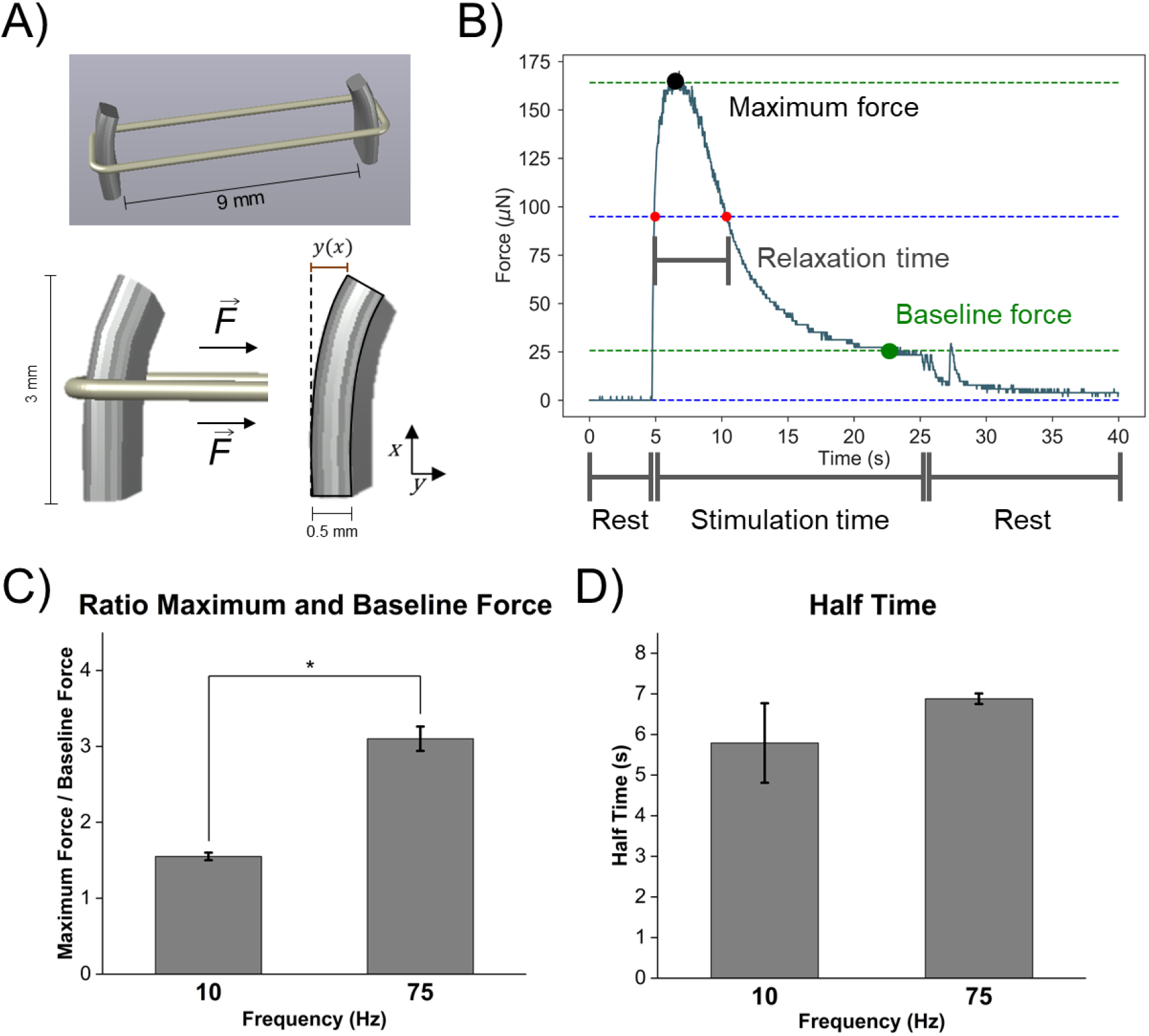
A) Representation of the mechanism of force monitoring with the drug testing platform. B) The maximum force, baseline force and relaxation time are extracted from 30 s high frequency stimulations (10 Hz or 75 Hz) in order to study differences in contraction kinematics. C) The ratio of maximum to baseline force indicate that 75 Hz stimulation produces a higher force stroke (N=4; mean ± standard error of the mean). D) The relaxation half time, however, is not affected by the frequency (N=4; mean ± standard error of the mean). * *p* < 0.05 (Student’s t-test).

Due to the interesting contraction profiles reported in Figure 2, we focused on the study of the kinetics and force generation of a high-(75 Hz) and a low-frequency stimulation (10 Hz) for 20 s. We extracted three main parameters from the contraction profiles: the maximum force achieved, the baseline force maintained until stimulation was stopped, and the relaxation time, defined as the time until achieving the median force between maximum and baseline after the onset of stimulation (Figure 4B). Initial observations reported differences in the maximum/baseline force ratio for 10 Hz and 75 Hz, confirming that higher frequencies produced a stronger initial twitch, as already reported^20,25^ (Figure 4C). The relaxation time, however, did not significantly differ between both frequencies (Figure 4D).

These contraction parameters were monitored before and after the addition of 40 ng/mL of TNF-α in order to test functional changes compatible with skeletal tissue aging. **Figure 5** shows these results for both frequencies, as well as control (untreated) and TNF-α-treated samples, before (0 h) and after (24 h) treatment with the cytokine. All contraction parameters for each sample were normalized to the average value in the pre-treatment state (0 h) in order to reduce initial variability common in replicas of biological studies. Control samples, in general, did not show significant changes in the contraction parameters, although there was high inherent sample variability, despite standardizing culture conditions by fixing exact concentration of cells and always using the same passage (Figure 5B). Samples treated with TNF-α, however, showed a marked decrease in maximum and baseline forces after 24 h of treatment, reaching half of their pre-treatment values (Figure 5C). This was statistically significant for the 75 Hz stimulation, although the same tendency was observed for 10 Hz, with higher variability. The graphs at the right-hand side compare the ratios of control and TNF-α-treated samples at 24 h. It can be seen how the maximum and baseline forces were also reduced in TNF-α compared to controls, therefore eliminating temporal effects related to the natural maturation of the tissue. Moreover, there was a slight tendency to increase the relaxation time after TNF-α treatment, although not significantly. The comparison of these ratios at 24 h (right-hand side plots) revealed that the addition of TNF-α seemed to increase the relaxation time of the hSMTs when compared to control samples. Finally, Figure 5C and Supplementary Videos S3 and S4 show representative examples of the contraction profiles for 75 Hz of samples with and without TNF-α after 24 h. Here, it can be seen how the cytokine TNF-α affects both the strength of the fast twitching response and the baseline tetanic force in the short term (24 h), and also slightly increases its relaxation time. This is consistent with previous studies, which show that muscle aging is associated to an increase in relaxation time and a decrease in the force output^20,33–35^. Thus, besides the morphological changes caused by TNF-α previously discussed, applying a high-frequency 75 Hz stimulation to TNF-α-treated tissues could resemble muscle dysfunctional changes that occur during aging, with typical dysfunctional changes, in particular loss of force

**Figure 5.**
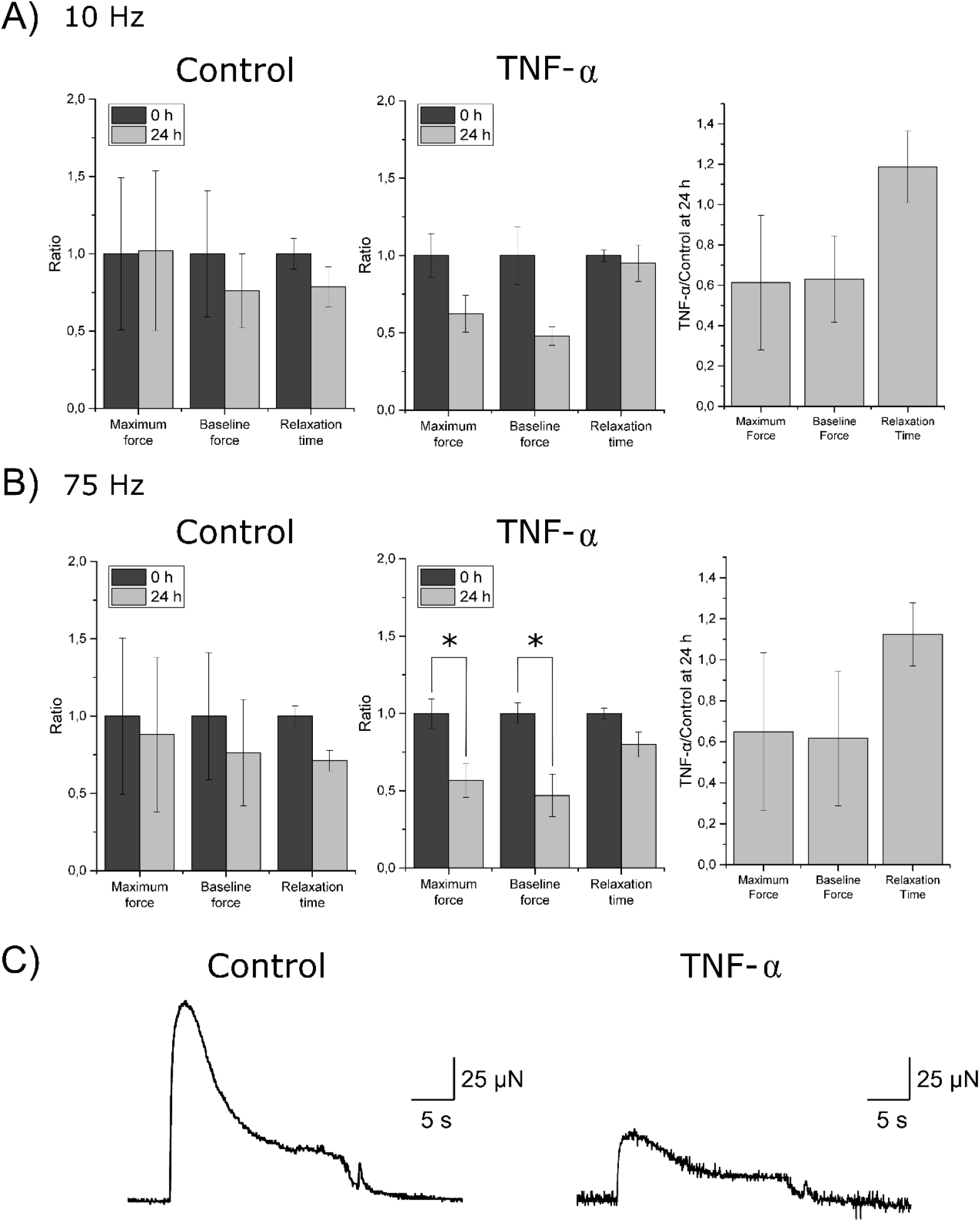
Effect of TNF-α on the contraction parameters previously defined during a sustained stimulation of A) 10 Hz and B) 75 Hz, both for control and TNF-α-treated samples. On the right side, the ratio of each parameter for TNF-α vs control at 24 h are compared to better see the differences between models. All data are normalized with respect to the initial conditions before any treatment (0 h). C) Representative example of the contraction profiles (75 Hz) for a control and TNF-α with the same scaling, to better appreciate the differences produced. N=3-6. All results are shown with mean ± standard error of the mean. * *p* < 0.05 (Student’s t-test).

To further exploit the possibilities of our drug testing platform, we tested the effects of Argireline^®^ Amplified peptide. As a first approach, the peptide was added to healthy 3D muscle models for 48 h, with a replenishment after 24 h at a concentration of 2 mg/mL. **Figure 6** shows the impacts on the contraction parameters after 24 h and 48 h of addition to non-aged muscle, where all parameters were normalized to the average value in the p re-treatment state (0 h). The results point at a frequency-dependent effect. For low frequencies (10 Hz), the maximum twitch force was increased after 48 h, while it was decreased for high frequencies (75 Hz). This decrease was especially prominent in the baseline force, which was radically reduced until reaching total inhibition of tetanic contraction for 75 Hz (see Supplementary Video S5). For 10 Hz, however, baseline force was not significantly changed, although slightly increased. The relaxation time was reduced for both frequency conditions, although more strongly for 75 Hz after 48 h.

**Figure 6.**
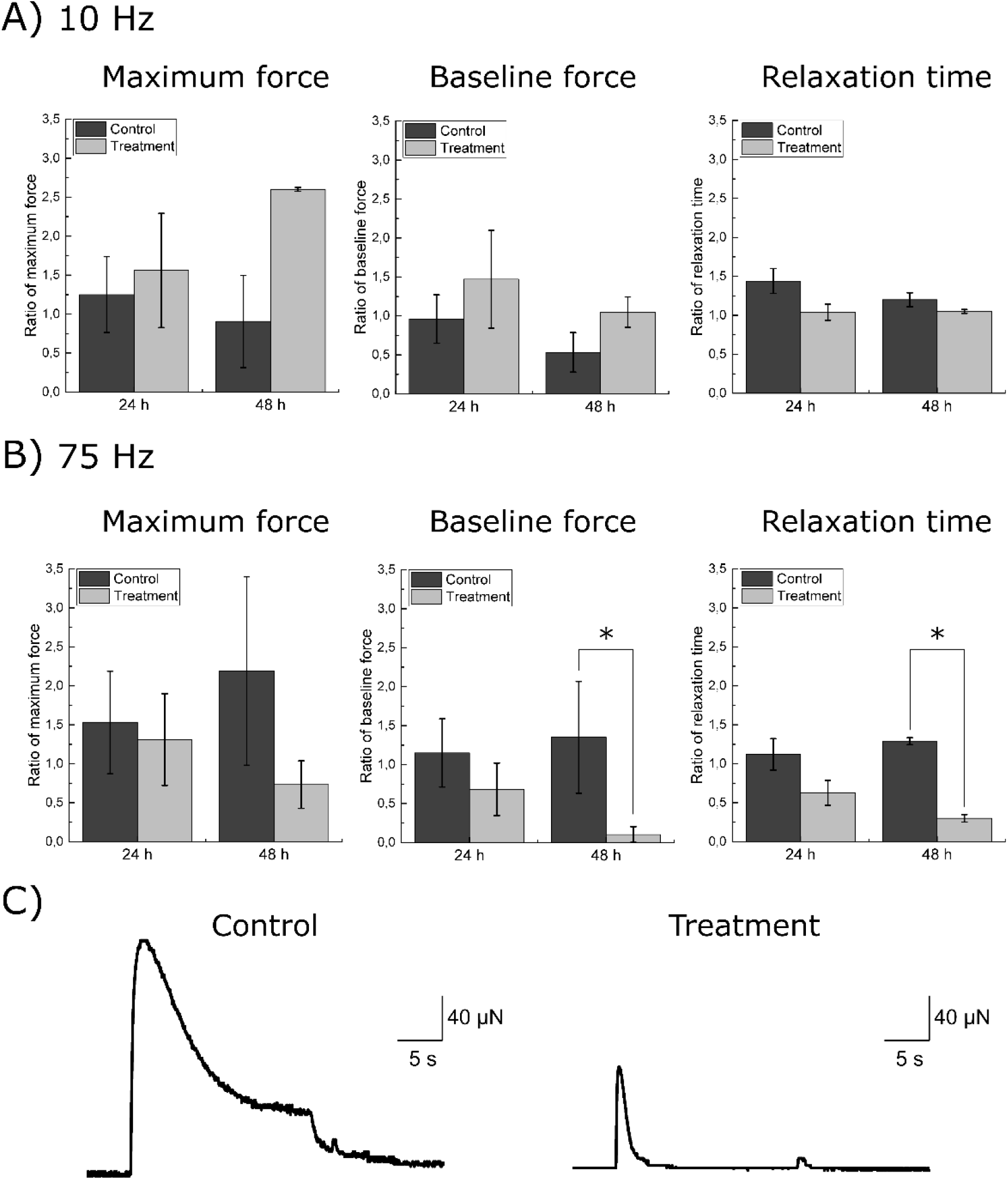
A) Effect of the Argireline^®^ Amplified peptide on the contraction parameters defined in figure 4 in a sustained stimulation of A) 10 Hz and B) 75 Hz, both for control and peptide-treated samples for 48 h. All data are normalized with respect to the initial conditions before any treatment (0 h). C) Representative example of the contraction profiles (75 Hz) for a control and a sample treated with the peptide with the same scaling, to better appreciate the differences produced. N=3-6. All results are shown with mean ± standard error of the mean. * *p* < 0.05 (Student’s t-test).

Representative contraction profiles of a 75 Hz stimulation can be seen in Figure 6C. It can be readily observed how, after 48 h, the tetanic contraction was partially inhibited despite having a continuous electrical stimulation. Only a short-lived twitch contraction remained, which decreased rapidly to a relaxed state. Therefore, the Argireline^®^ Amplified peptide showed relaxing effects by inhibiting maximum and baseline forces, and also reducing relaxation time in young/healthy muscles, leading to an inhibition of sustained contractions upon high frequency stimulations (75 Hz).

The effects of this peptide were also studied on a model of aged hSMT, also revealing frequency-dependent responses (**Figure 7**). I n this study, TNF-α was introduced in the culture media for 24 h in order to induce the aging attributes revealed in the previous figures (Figure 7A). Here, this point was considered the initial phase for this study (0 h), and all contraction parameters were normalized to the average value of the corresponding samples at this stage. Subsequently, the samples were treated with the peptide for two additional days (time-points 24 h and 48 h). For 10 Hz stimulations, both the maximum twitch and baseline forces increased when the peptide was added, in contraposition with samples to which no treatment was applied, similar to the results obtained for the same frequency in Figure 6. The relaxation time, however, did not show considerable changes after 48 h of peptide treatment. At 75 Hz, the condition previously described to mimic muscle physiological state during aging, peptide treatment lead to a decrease in the maximum force generation after 48 h. Likewise, the baseline force decreased significantly after 48 h of treatment with the peptide, as reported in Figure 6, although it did not reach a complete inhibition of the contraction. The relaxation time was modestly decreased after addition of the peptide, as also shown in Figure 6. Considering these results, our data seem to indicate that treatment with the peptide at 75 Hz can reduce maximal force and provides faster relaxation in both young and aged-induced muscle, which could provide cosmetic benefits to fight against skin aging appearance.

**Figure 7.**
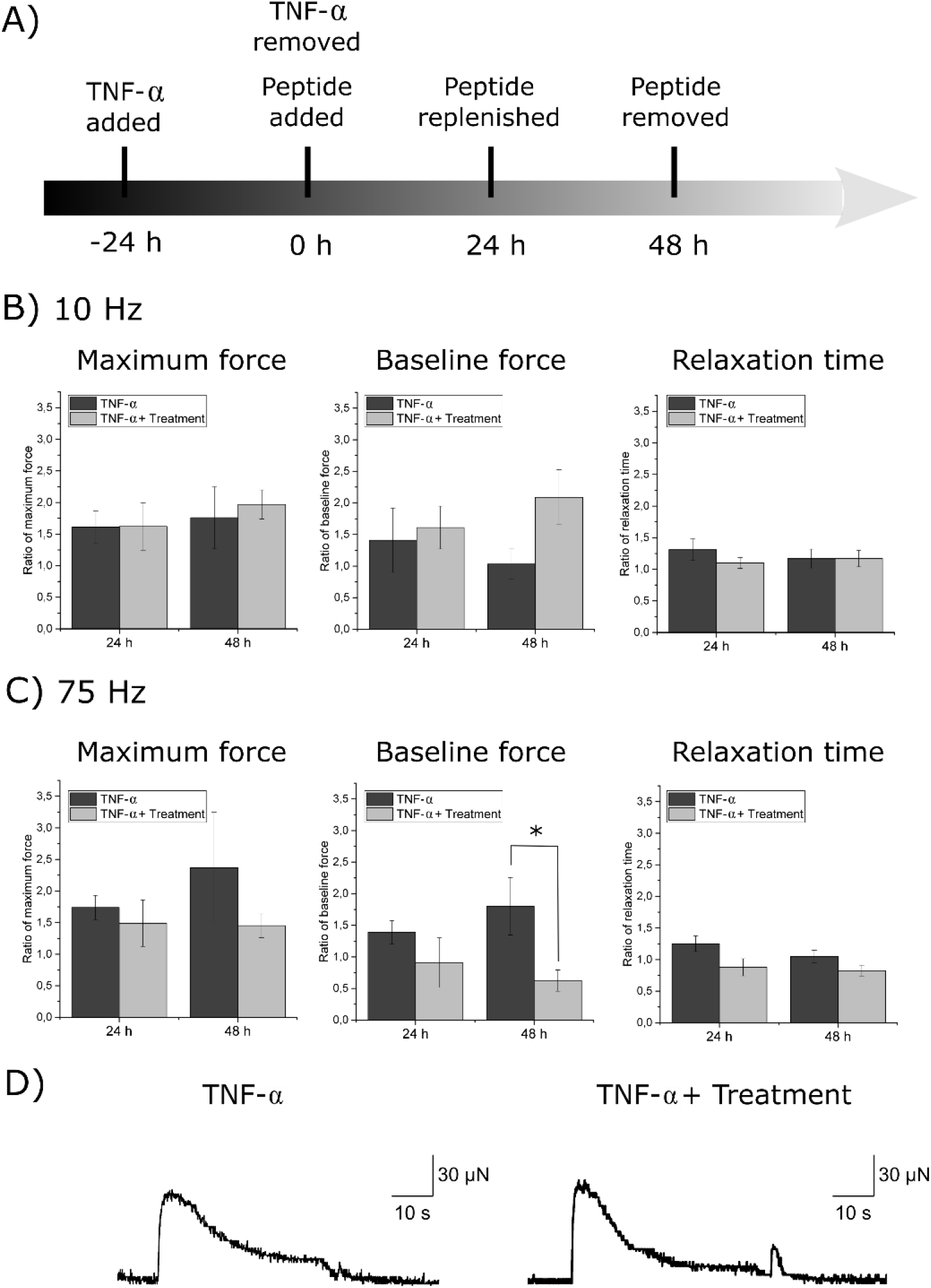
A) Combined effect of TNF-α and the Argireline^®^ Amplified peptide in the contraction parameters defined in Figure 4 in a sustained stimulation of B) 10 Hz and C) 75 Hz. All data are normalized with respect to the initial conditions of aged muscle before the treatment with the peptide (0 h). C) Representative example of the contraction profiles (75 Hz) for a sample treated with TNF-α and a sample treated with TNF-α and the peptide with the same scaling, to better appreciate the differences produced. N=3-6. All results are shown with mean ± standard error of the mean. * *p* < 0.05 (Student’s t-test).

## 3. Conclusions

In this work, we presented a 3D-printed and versatile platform for the study of young and aged human muscle functionality and its potential applications as a drug testing platform. We 3D-bioengineered a model of aged or senescence-like hSMT by the addition of TNF-α, and we showed its effects both in morphology (by immunostaining) and functionality (by contraction profiles), characterized by a reduction in fiber diameter and loss of nuclei, as well as loss of maximum and baseline forces. To further illustrate the possibilities of the testing platform, we tested, as a proof of concept, the impacts of Argireline^®^ Amplified peptide in the contraction kinetics of bioengineered human muscle, showing model-dependent effects (aged *vs* non-aged). We believe that this drug-testing platform could have a potentially positive impact for the development of new drugs in the biomedical or cosmetic fields, as it can mimic the 3D environment of native hSMT, as well as extract useful information of all kinds of stimulation profiles, both acute and chronic. Moreover, we are confident that our work could provide valuable results towards the development of more relevant 3D models of adult and aged skeletal tissue, as the effects that we observed after addition of TNF-α coincide with those reported in the literature. Nevertheless, better and more advanced models for aging could include satellite cells, which seem to be especially affected by TNF-α^26^. Towards more specialized treatment, human myoblasts from patients suffering from rare muscular disorders, such as muscular dystrophies, could be used to understand their development and treatment.

## 4. Materials and methods

### 4.1. Cell culture

Purchased cryopreserved Human Skeletal Myoblasts were obtained from the abdominal area of a Caucasian female donor aged 28 (Zenbio, # cat. SKB-F-1). Their growth medium was Skeletal Muscle Cell Growth Medium (PromoCell, # cat. C-23060) supplemented with Skeletal Muscle cell Growth Medium SupplementMix (PromoCell, # cat. C-39365) and differentiation medium was Skeletal Muscle Cell Differentiation Medium (PromoCell, # cat. C-23061) supplemented with Skeletal Muscle Cell Differentiation Medium SummplementMix (PromoCell, # cat. C-39366). Cryovials were thawed and 2D cultures were expanded and passaged until reaching passage 4. Once the cells reached a confluence of 50-70% (approximately after 5-6 days growth) they were recollected. In this process, T-75 flasks were rinsed twice with PBS 1x and 3 mL of trypsin were added to each flask for 2 min. After cells detached, trypsin was neutralized with 9 mL of supplemented Skeletal Muscle Cell Growth Medium and final volume was centrifuged at 300 x g for 5 min. Supernatant was discarded and the pellet was resuspended in supplemented Skeletal Muscle Cell Growth Medium for cell counting. Eventually, the resuspended pellet was once again centrifuged at 300 x g for 5 min and the resulting pellet used for the sample 3D fabrication.

### 4.2. 3D printing of molds and posts

In order to fabricate the PDMS molds for cell-laden hydrogel casting and posts for the force measurement platform, the 3D bioprinter INKREDIBLE+ from CELLINK® was used. Both structures were made of polydimethylsiloxane (PDMS; Dow Corning SE 1700) with a crosslinker ratio of 1:20 (coming with the base material as a kit), mixed according to the manufacturer’s instructions. The designs of the structures were carried out in AutoCAD (v. 2019), exported as .stl files, and transformed into GCode to be 3D printed at a pressure of 220 kPa. After printing, the molds were cured at 80 °C for 16 h and posts were cured at 37 °C for 72 h. The walls of the molds were 1 mm high and their inner and outer diameters were 10 mm and 13 mm, respectively. The posts were 3 mm high, 0.5 mm wide and with 2 mm of lateral width, separated by a distance of 9 mm.

### 4.3. 3D sample fabrication

For the fabrication of the hydrogel, a mixture of supplemented Skeletal Muscle Cell Growth Medium (16.25% v/v), 50 U/mL of thrombin (3.75% v/v), Matrigel^®^ (30.0% v/v) and fibrinogen 8 mg/mL (50.0% v/v or 4 mg/mL final concentration) were used. These concentrations were chosen according to previous reports in the literature that optimized this mixture^14^, in particular Hinds et al., where they found that the range of 20-40% of Matrigel^®^ and 4-6 mg/mL of fibrinogen gave optimal force output^43^. First, cultured human skeletal myoblasts at passage 4 and reaching 80% confluency were detached with trypsin as previously described. The resulting cell pellet after centrifugation was mixed with all the components except fibrinogen at a temperature in a cold ice bath, when Matrigel® is fluid. Once the solution was homogenized, fibrinogen was added and then, 80 µL of the final mixture was poured inside the mold, fast enough to avoid coagulation due to the fibrinogen to fibrin crosslinking. The density of cells in each mold was always 5 million/mL. After this, molds were incubated for 30-60 min to allow Matrigel to crosslink. Then, 1.5 mL of Skeletal Muscle Cell Growth Medium with 1 mg/mL 6-aminocaproic acid (ACA) was added to the Petri dish. Samples were put into the incubator for 48 h. At 24 h samples were gently lifted from the outer wall of the mold to enhance tissue compaction.

After 48 h samples were entirely lifted from the mold and carefully transferred to Petri dishes containing the 3D printed posts. The ring of cells should encircle both pillars. Supplemented Skeletal Muscle Cell Differentiation Medium was added, and the samples were put into the incubator until they have reached optimum compaction and were able to contract under electrical stimulation. During this time, medium was changed every 38-72 h.

### 4.4. Electrical pulse stimulation

The setup for the electrical pulse stimulation was composed by a waveform generator (PM8572, Tabor Electronics), a signal amplifier x15, an oscilloscope (DS1104Z, Rigol) and a set of handmade electrodes based of graphite rods (cat. number 30250, Ladd Research) separated by a distance of 12.5 mm. The electrodes are made of two cylindric graphite rods (cat. number 30250, Ladd Research) placed on opposite sides of the Petri dish. The recording was carried out inside an inverted microscope (DMi8, Leica) and samples are placed in a chamber that allows to mimic physiological conditions (37 °C and 5% CO2). Pulses of different frequencies (depending on experiment) of 2 ms were applied, keeping a constant electric field of 1 V/mm between the electrodes. Full characterization of this electrodes can be found in the Supplementary Information of a previous publication^2^.

### 4.5. Measurement of force and contraction kinetics

The force that 3D fabricated samples exert against the pillars is measured by the displacement of the pillars normalized by the height of the pillar at which the sample is applying the force under a certain voltage and frequency conditions. For this, the Z-stack option of the Leica DMi8 microscope was used to calculate the distance from which the tissue was pulling. Then, videos were recorded at 50 FPS and a homemade Python code was applied to recognize pixel displacement. This code was designed to detect the edges of the different structures by applying a thresholding based on a canny edge detection algorithm. The user manually adjusted the threshold if necessary and selected a straight line in the border of the pillar which indicated the position along which the displacement detection was being analyzed. Eventually, the displacement of the pillars during the whole recorded video was processed and force was calculated according to a beam bending theory, as previously reported^2^, using the equation 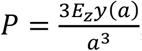, where *P* is the applied force, *E* is the Young’s modulus, *I*_*z*_ is the second moment of area of the post around the z axis, *a* is the height from which the tissue is pulling and *y*(*a*) is the displacement of the post at such height. The Young’s modulus of the PDMS was measured to be *E* = 206 kPa, calculated by fitting the linear range of 15-30% strain after a mechanical test, as previously reported^2^. For contraction kinetics initially studied in Figure 2, a home-made Python script based on a tracking algorithm was applied. A region of the tissue was manually selected, and its position tracked throughout the video. Then, the image distance between the tracked regions in time was computed to obtain the contraction kinetics in arbitrary units.

### 4.6. Calcium imaging

Ca-imaging was performed in an inverted microscope at 37 °C and 5% CO2 atmosphere. hSMTs were incubated with a calcium-sensitive Fluo-4 dye (F10489, ThermoFisher) and stimulated as previously described. The extracted videos were analyzed with a home-made Python script in which the user selected a region of interest and the total pixel intensity in that region was plotted as a function of time.

### 4.7. Tumor Necrosis Factor α and peptide treatment

Some samples are treated with HumanKine™ Tumor Necrosis Factor α (TNF-α), human recombinant, expressed in HEK 293 cells (Sigma-Aldrich, # cat. H8916). The cytokine was added at different concentrations (20 ng/mL, 40 ng/mL or 80 ng/mL) in the culture media for 24 h. Argireline^®^ Amplified peptide obtained from Lipotec™ Active Ingredients and added, when corresponding, at a concentration 2 mg/mL in the culture media for 24 h. Then, it was replenished for an additional 24 h.

### 4.8. Live/dead and immunostaining

Different types of analysis are performed in order to extract information about the viability of the 3D fabrication method and the correct differentiation and maturation of the tissue. Cell viability was assessed by PrestoBlue*™* cell viability reagent was purchased from Thermo Scientific™ (A13262) and used according to the manufacturer’s instructions. Before replenishing culture media at D0, D2 and D5 of differentiation, PrestoBlue*™* reagent was added at a ratio of 1:10 with respect to the total volume of the dish. After 2 h of incubation at 37 °C, 10 μL of media was taken and its absorbance measured with a Spark^®^ multimode microplate reader at 560 nm, normalized to the value at 600 nm, subtracting the background of only-medium controls. For the visualization of differentiated myoblasts, samples are immunostained using Myosin 4 Monoclonal Antibody (MF20) conjugated with Alexa Fluor 488 (eBioscience™) and Hoechst staining to label myosin and nuclei, respectively. ImageJ/Fiji software is used to process the results.

### 4.9 RT-qPCR

Total RNA content of 3-4 biological replicates of control and samples treated with TNF-α at 40 ng/mL for 24 h were extracted using the RNeasy mini kit (QIAGEN, 74134) according to manufacturer’s instructions. The purity and concentratio of the extracted RNA was assessed with with Nanodrop ND-1000 (Thermo Scientific™). A total amount of 500 ng of RNA was converted into cDNA using the ReverAid First Strand cDNA Synthesis Kit (Thermo Scientific™, K1622). RT-qPCR reactions were performed with PowerUp SYBR Green Master Mix (Applied Biosystems, A25742), according to manufacturer’s instructions, with 500 ng of cDNA and the target primers in a total volue of 10 μL in a StepOnePlus Real-Time PCR System (Applied Biosystems, 4376600). The target genes were normalized to the expression of GAPDH. Melt-curve tests were performed to ensure that only one amplicon was being amplified. Negative and non-templated controls were also performed to ensure purity of the samples and reagents. The following primers were used:

- GAPDH: FW (5’ CCTGACCTGCCGTCTAGAAA 3’) RV (5’ TGCTGTAGCCAAATTCGTTG 3’)
- Atrogin-1: FW (5’ GCAGCTGAACAACATTCAGATCAC 3’) RV (5’ CAGCCTCTGCATGATGTTCAGT 3’)
- MuRF-1: FW (5’ CCTGAGAGCCATTGACTTTGG 3’) RV (5’ CTTCCCTTCTGTGGACTCTTCCT 3’)
- LC3b: FW (5’ GAGAAGCAGCTTCCTGTTCTGG 3’) RV (5’ GTGTCCGTTCACCAACAGGAAG 3’)
- MyHCI type 2X: FW (5’ CTGGCTCTCCTCTTTGTTGG 3’) RV (5’ GAGAGCAGACACAGTCTGGAAA 3’)
- MyHCII type 2A: FW (5’ GGAGAGGGAGCTGGTGGA 3’) RV (5’ CTCTCTGAAAAGGGCAGACA 3’)
- MyHC8 neonatal type: FW (5’ TCTTCTGGAAGAAATGAGAGATGA 3’) RV (5’ GCTTCTCTCCTTTGCAACATC 3’).

### 4.10. Statistical data treatment

The statistical test used to perform multiple data comparison between myocyte diameter and number of nuclei per fiber (Figure 2) was one-way ANOVA followed by post-hoc analysis (Tukey’s Multiple Comparison test; N=12). Differences in expression of genes with RT-qPCR were assessed with a Student’s t-test. Normality was assessed with the Shapiro-Wilk test and homoscedasticity with the F-test test in Prism (v. 8.3.1). When the conditions for a t-test were not fulfilled, a Mann-Whitney test was performed. In any case, the only gene expression that showed significant differences, MuRF-1, fulfilled both conditions of normality and homoscedasticity.

To analyze the difference in contraction parameters at different frequencies between two group (Figure 3), a Student’s t-test was performed (N=4). Homoscedasticity was checked by the normal Q-Q plot and the equality of variances by the residuals versus fit plot in Python (v. 3.7). For the remaining experiments, data from the different contraction parameters were collected for each sample at different frequencies and for several days, as explained in the results section. For each of the three figures reporting comparisons between these data (Figures 5, 6 and 7), the following data treatment was applied. For Figure 5, whose aim was to detect differences after the addition of TNF-α after 24 h, each value of maximum force, baseline force or relaxation time was normalized to the average value within its sample group (control or TNF-α) at 0 h (before adding the cytokine). In this way, we wanted to reduce the initial variability in replicates, since we were only interested in the time evolution of the samples. A t-test of dependent (paired) samples was used to detect statistical differences between samples at 0 h and 24 h. A variance test (F-test) was performed beforehand to check whether we could account for equal variances or not in the t-test and the Kolmogorov-Smirnov (K-S) test to check for normality. For Figure 6, whose aim was to detect differences between control and peptide-treated groups at different time-points, all contractions parameters were also normalized to the average value for each group at 0 h (before adding the peptide). To test for significance, an independent-sample t-test was performed, also preceded by a variance F-test and normality K-S test. Finally, for Figure 7, whose aim was to detect differences in the contraction parameters when the peptide was added in aged muscle models, the values were normalized to their group average at 0 h, the time-point when TNF-α was removed and the peptide not added yet (see time-line in Figure 7A). As in the previous case, an independent-sample t-test was performed to check for statistical differences, preceded by a variance F-test to confirm or not variance equality and a normality K-S test. Statistical tests were carried out in R. The level of significance was set to *p*-value ≤ 0.05. For every experiment of Figures 5, 6 and 7, the population was N = 3-6.

## ACKNOWLEDGEMENT

R.M. thanks “la Caixa” Foundation through IBEC International PhD Programme “la Caixa” Severo Ochoa fellowships (code LCF/BQ/SO16/52270018). T.P. thanks the European Union’s Horizon 2020 research and innovation program, under the Marie Sklodowska-Curie Individual Fellowship (H2020-MSCA-IF-2018, DNA-bots). M.G. thanks MINECO for the Juan de la Cierva fellowship (IJCI2016-30451), the Beatriu de Pinós Programme (2018-BP-00305) and the Ministry of Business and Knowledge of the Government of Catalonia. S.S. acknowledges the CERCA program by the Generalitat de Catalunya, the Secretaria d’Universitats i Recerca del Departament d’Empresa i Coneixement de la Generalitat de Catalunya through the project 2017 SGR 1148 and Ministerio de Ciencia, Innovación y Universidades (MCIU) / Agencia Estatal de Investigación (AEI) / Fondo Europeo de Desarrollo Regional (FEDER, UE) through the project RTI2018-098164-B-I00. This project was also partially funded by Agencia Estatal de Investigación (CEX2018-000789-S). The authors thank the Lipotec™ Active Ingredients’ team for developing and providing with Argireline® Amplified peptide and the staff at IBEC’s MicroFabSpace for the technical support in CSLM experiments.

## ABBREVIATIONS

hSMT: human skeletal muscle tissue
TNF-α: tumor necrosis factor α
GM: growth medium
DM: differentiation medium
ACA: 6-aminocaproic acid
PDMS: polydimethylsiloxane

