## Supplementary Information for "3D-printed drug testing platform based on a 3D model of aged human skeletal muscle"

### Supplementary Videos

**Video S1 – Stimulation at 1 Hz**. Video showing a hSMT fiber electrically stimulated at 1 Hz for a couple of seconds (starting at 5 s).

**Video S2 – Calcium imaging**. Calcium imaging of a hSMT fiber electrically stimulated at 1 Hz for some seconds.

**Video S3 – Control sample after 24 h - 75 Hz stimulation**. Video showing the contraction of the tissue, along with the force measurement, of a control muscle fiber electrically stimulated at 75 Hz.

**Video S4 – TNF-a sample after 24 h - 75 Hz stimulation**. Video showing the contraction of the tissue, along with the force measurement, of a muscle fiber treated with TNF-α for 24 h, electrically stimulated at 75 Hz.

**Video S5 – Peptide-treated sample after 48 h - 75 Hz stimulation**. Video showing the contraction of the tissue, along with the force measurement, of a muscle fiber treated with the peptide for 48 h, electrically stimulated at 75 Hz.

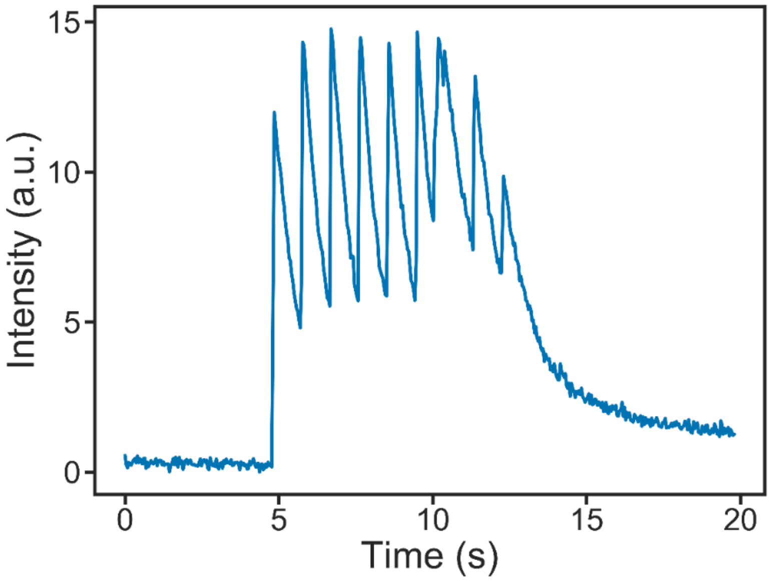


**Figure S1.** Calcium imaging of a sample stimulated at 1 Hz for
